# DNA-BOT: A low-cost, automated DNA assembly platform for synthetic biology

**DOI:** 10.1101/832139

**Authors:** Marko Storch, Matthew C. Haines, Geoff S. Baldwin

## Abstract

Multi-part DNA assembly is the physical starting point for many projects in Synthetic and Molecular Biology. The ability to explore a genetic design space by building extensive libraries of DNA constructs is essential for creating programmed biological systems that perform the desired functions. With multiple DNA assembly methods and standards adopted in the Synthetic Biology community, automation of the DNA assembly process has received serious attention in recent years. Importantly, automating DNA assembly enables larger builds using less researcher time, increasing the accessible design space. However, these benefits currently incur high costs for both equipment and consumables. Here, we address this limitation by introducing low-cost DNA assembly with BASIC on OpenTrons (DNA-BOT). For this purpose, we developed an open-source software package dnabot (https://github.com/BASIC-DNA-ASSEMBLY/dnabot). We demonstrate the performance of DNA-BOT by simultaneously assembling 88 constructs composed of 10 genetic parts, exploring the promoter, ribosome binding site (RBS) and gene order design space for a 3-gene operon. All 88 constructs were assembled with high accuracy, at a cost of $1.50 - $5.50 per construct. This illustrates the efficiency, accuracy and affordability of DNA-BOT making it accessible for most labs and democratising automated DNA assembly.

## Introduction

Creating DNA constructs is the foundational process that allows biologists to interrogate and engineer biological systems for a wide range of applications in basic research, biotechnology and more recently data storage^1,2^. Consequently, DNA assembly techniques and standards have evolved to address the desire to construct diverse sequences ranging in sizes from plasmids to whole genomes^1^. As with many routine molecular biology methods, workflow standardisation has enabled DNA assembly techniques to be completely automated, increasing the scale of construction, extending the addressable design space. This approach has now led to the emergence of Biofoundries^3,4^.

We previously developed the Biopart Assembly Standard for Idempotent Cloning (BASIC) method and standard to enable highly accurate multi-part DNA assembly at both manual bench and fully automated Biofoundry scale^5,6^. BASIC defines a single biopart storage format with prefix and suffix sequences flanking each part and provides a method for highly accurate linker-based DNA assembly. Further, BASIC enables idempotent cloning where each assembled cassette becomes a BASIC biopart, ready for subsequent hierarchical assembly through the exact same automation-friendly workflow. For convenience, and with automation in mind, a collection of neutral and functional linkers encoding RBS sequences or fusion peptides was designed and is available from Biolegio in a ready to use low-cost consumable in 96-well plate format^7^. Automated cloning performance depends strongly on assembly accuracy and efficiency to minimize the number of clones that need to be screened. BASIC provides this high accuracy and efficiency through long linker overhangs, guiding the assembly of linker-ligated parts. Currently, BASIC and alternative DNA assembly methods have only been automated using expensive infrastructure, limiting community access to the benefits automated DNA assembly brings to research and applications in biology^6,8–11^.

With the recent advent of the Opentrons OT-2 liquid handling robot, equipment costs for entry-level automation dropped significantly, making it accessible to most molecular biology researchers^12^. The OT-2 can accurately transfer volumes from 1 to 300 μL with single or 8-channel pipettes and supports the BASIC integration through additional modules for automated temperature control and magnetic bead manipulation. A further advantage is the open-source, python-based application programming interface that facilitates rapid protocol development.

Here, we present the DNA-BOT platform, which combines highly accurate, open source BASIC DNA assembly with the low-cost Opentrons OT-2 for automated DNA assembly. We hypothesised DNA-BOT would be affordable for most research groups, while achieving the accuracy needed for large-scale, automated projects. Not only would this add to the available SynBioStack, it would improve the community’s ability to iterate through Design-Build-Test-Learn cycles, driving the development of Synthetic Biology^13^.

## Results and discussion

BASIC DNA assembly is performed in 4 separate steps (Figure 1a). These were implemented as four individual processes on the OT-2, each with a dedicated deck setup (Figure S1) for the associated script (Figure 1b). Briefly, in the first step BASIC clips are created by digesting BASIC parts out of their storage vectors and simultaneously ligating linkers that define the assembly order, in a one-pot enzymatic reaction (**Step 1**). The resulting clips are purified from un-ligated linkers using solid-phase reversible immobilization (SPRI) paramagnetic beads (**Step 2**). These purified clips expose ~20 base overhangs, facilitating their assembly when incubated at an appropriate temperature in annealing buffer. (**Step 3**). Subsequent transformation of assembled constructs and plating on selection media (**Step 4**) yields colonies for downstream assays and applications.

**Figure 1.**
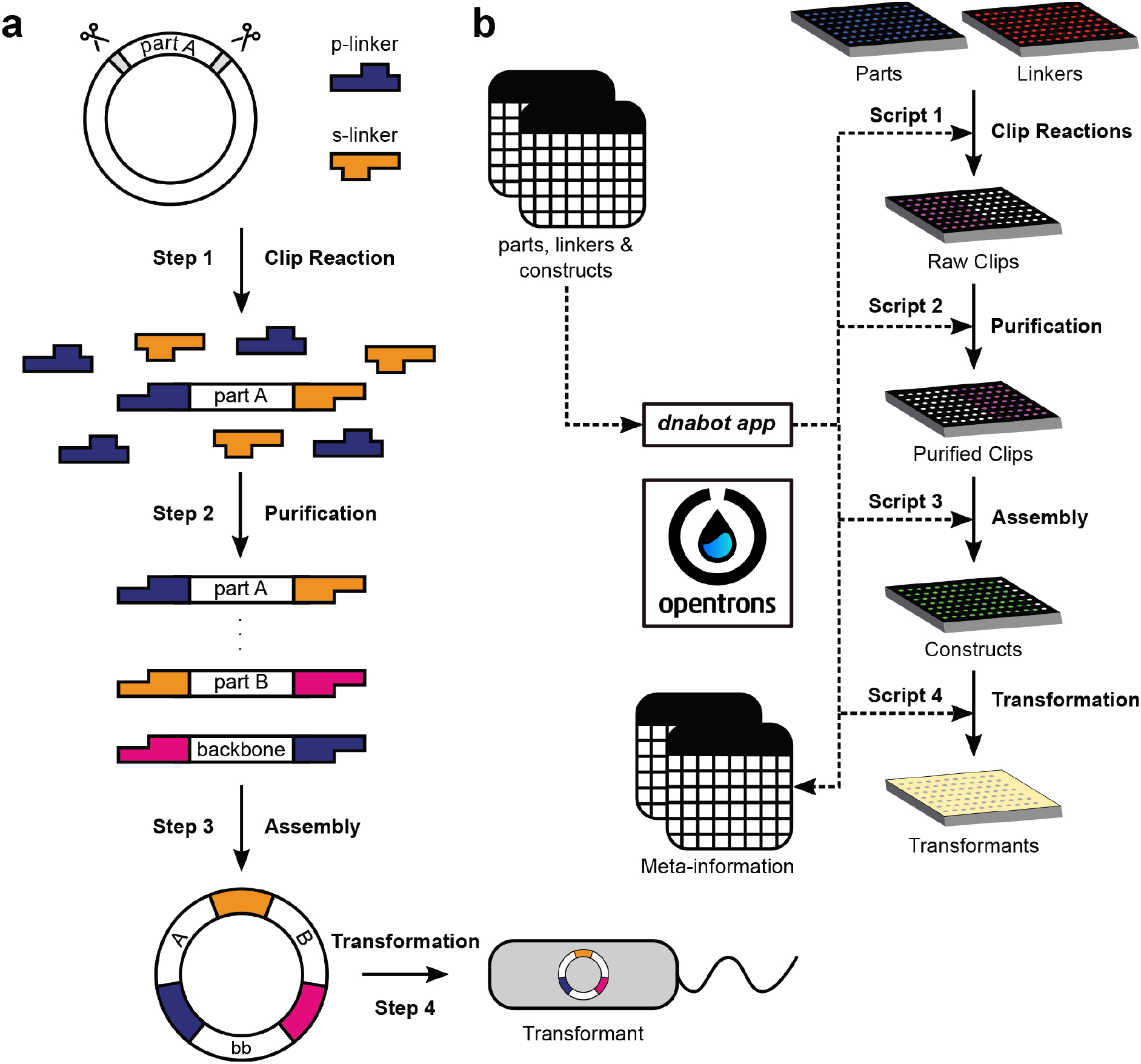
BASIC DNA assembly and the DNA-BOT workflow. (a) BASIC DNA assembly workflow: Step 1 - Clip Reaction: Simultaneous digestion and ligation attaches prefix (p-) and suffix (s-) linker sections to parts. Step 2 - Purification: Clips are purified from raw clip reactions via Solid Phase Reversible Immobilization (SPRI), removing excess linkers and enzymes. Step 3 - Assembly: Purified clips are annealed, forming circular constructs e.g. part A & B are annealed with a backbone (bb) part in a 3-part assembly. Step 4 - Transformation: Assembled constructs are transformed into *E. coli*. (b) csv files describing source plates for parts, linkers and construct designs are processed by the dnabot application (app), returning OT-2 scripts along with meta-information csv files. Each script runs the corresponding BASIC step in microtiter plate format, finally spotting colonies on selective LB-agar plates. Dotted and solid-lines denote information flow and OT-2 runs, respectively.

After developing the principles of the robotic protocols to implement the four BASIC steps, we created an open-source python package (https://github.com/BASIC-DNA-ASSEMBLY/dnabot) that provides a convenient interface to generate scripts and associated parameters for the assembly and transformation of up to 96 constructs using BASIC parts and Biolegio BASIC Linkers^7,14^. The dnabot application reads csv files detailing construct designs and plates containing BASIC parts and linkers to be used in a given project. Following the acquisition of these parameters the designs are analysed and parsed into the required clip reactions and assembly instructions, directing the generation of four specific OT-2 scripts along with associated meta-information (Figure 1b).

During Step 1, up to 48 clip reactions are setup, before user-mediated transfer to an external thermocycler, providing the clip reaction conditions (**Script 1**). In Step 2, the OT-2 magdeck module is used to purify raw clip reactions from the left half of the 96-well plate using SPRI beads, depositing purified clips in the right half of the plate (**Script 2**). In Step 3, purified clips are combined in annealing buffer to assemble specified constructs by annealing in an external thermocycler (**Script 3**). In Step 4, assembled constructs are mixed with competent cells on the OT-2 before heat-shock transformation using an external thermocycler. After recovery in SOC medium, liquid cultures of transformed cells are spotted on a selective LB-agar plate (**Script 4**). Script 4 takes advantage of the OT-2 temperature deck which enables transformation set-up and outgrowth at 4°C and 37°C, respectively. During the execution of these four scripts the Opentrons app will instruct the user to setup the OT-2 deck space as required (Figure S1), while prompting a few manual actions e.g. heat shock. Additionally, meta-information guides users through the composition of the required Clip Reaction Master Mix and the location of specific reagents. In the presented version DNA-BOT automates BASIC DNA assembly using only Opentrons equipment and an external thermocycler as hardware in standard lab settings (Supporting Information: DNA_BOT_instructions_v1.0).

To test DNA-BOT’s utility and ability to work at a relevant scale, we designed 88 constructs (Figure 2a) for assembly and transformation in parallel during a single run. Each variant encoded an operon expressing three fluorescent reporters GFP, RFP and BFP (green, red and blue fluorescent protein) on a p15A backbone with a Chloramphenicol-resistance cassette (Materials and methods). For these 88 constructs, 4 different promoters were used along with 2 or 3 different RBSs for each gene in 2 different gene orders. This design required 38 clip reactions to create the components for assembly of the final 88 constructs. In assembling these expression constructs, we benchmarked DNA-BOT’s performance while exploring an operon design space; one of many possible applications. Each construct consisted of 5 BASIC parts and 5 BASIC linkers with their identity defined for each variant in a construct design csv file. Together with csv files describing part and linker plates, the dnabot application generated the four required scripts and meta-information for assembly and transformation. All files are available at https://github.com/BASIC-DNA-ASSEMBLY/dnabot.

**Figure 2.**
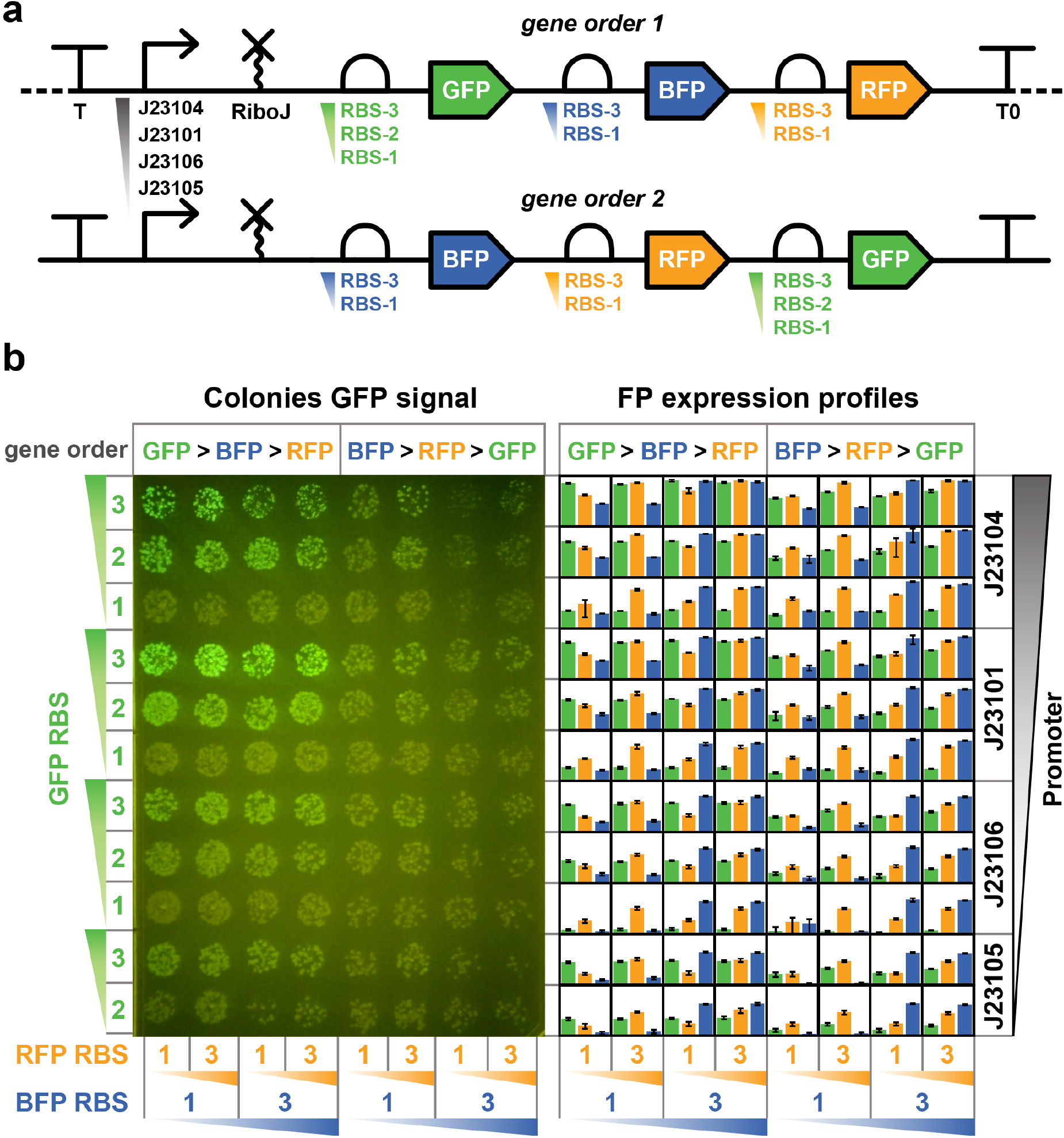
DNA-BOT provides for robust and accurate DNA assembly. Relative promoter & RBS strengths are indicated by gradients (a) SBOL Visual^18^ illustration of 88 constructs assembled using DNA-BOT. The library contained full permutations of promoters and RBSs as indicated in 2 gene orders. (b-left) Image of the agar plate on a Safe Imager™ 2.0 Blue Light Transilluminator, acquired following the DNA-BOT workflow with 10 μL of each transformation reaction spotted. Operon design features are indicated at axes. Green colonies indicate strong GFP expression. (b-right) Green, orange & blue bars denote normalised, mean GFP, RFP & BFP fluorescence, respectively, measured via flow cytometry for each construct from 3 biological repeats. Fluorescence is log scaled. Error bars denote standard deviations.

The workflow for the 88 assemblies was executed using the generated scripts and instructions, with the resulting transformants spotted in volumes of 5 and 10 μL onto SBS-LB-agar plates (Figure S2, materials and methods). Colonies were obtained for all 88 constructs and transformation control plasmids as expected (Figure 2b, Figure S2). The transformants were analysed for GFP fluorescence (Figure 2b-left) and triplicates for each assembly were picked for propagation in overnight liquid cultures. Cultures were analysed for GFP, RFP and BFP fluorescence at the single cell level via flow cytometry (Figure 2b, Figure S3, Materials and methods). These measurements enabled us to assess assembly success based on number and phenotypes of respective transformants across the 88 designs.

Observing the LB-agar plate in Figure 2b-left, we found each of the 88 spots contained a minimum of 5 colonies, returning transformants for all 88 constructs, illustrating DNA-BOT’s efficiency. Furthermore, cells exhibiting a pink phenotype were undetectable indicating a low background of un-digested backbone plasmid that would express the mScarlet counter-selection marker. Colonies within each spot show a largely homogeneous GFP expression phenotype as one would expect if they carried the same expression construct. From the flow cytometry of biological triplicates, we calculated and plotted mean and standard deviations derived from background corrected and normalised geometric means and arranged them in the corresponding agar plate layout in Figure 2b-right. The flow data demonstrates the different genetic designs led to a diverse range of fluorescence outputs ranging over 4-orders of magnitude (Figure S3). For each discrete design, we observed small standard deviations in the fluorescence response in almost all cases (Figure 2b-right and Supporting Information: DNA_BOT_flow_data).

From both the homogeneous GFP intensity of colonies within spots on the LB-agar plate and the small standard deviations observed from the flow cytometry measurements, we conclude that cells transformed with the same assembly have identical phenotypes, thus demonstrating that DNA-BOT provides high accuracy, in line with previous reports^5^. Furthermore, the trends observed in the expression profiles of the 3 fluorescent reporters reflect the expected positive correlations between promoter strength, RBS strength, proximity to the start of the operon and expression strength, typically governing gene expression within operons^15^. In conclusion DNA-BOT performs DNA assembly with high efficiency and high accuracy.

During this automated workflow the OT-2 performed 1578 pipetting steps, 38 magnetic bead purifications and 96 heat-shock transformations in 96-well format. The OT-2 including all required modules and pipettes costs around $8k and the cost per construct was estimated to be $ 1.50 or $ 5.50, depending on whether in-house or commercial competent cells were used, respectively (Table S2). We estimate the hands-on time for the whole process to be around 1 hr, 30 minutes (Table S3). This compares favourably with 4-5 hrs when implementing the same process manually. While this illustrates significant time saving, more importantly the process is more robust and reliable, since robots typically outperform humans in repetitive tasks e.g. cherry-picking liquid transfers of small volumes.

While the current performance of DNA-BOT is already very useful, we see several opportunities for future development. For instance, Opentrons will soon offer an onboard thermocycler for the OT-2. This will allow users to implement DNA-BOT relying on low-cost Opentrons hardware only^12^. Next, we will continue to develop the DNA-BOT software package to integrate with open-source DNA design tools like SBOL^16–18^ improving UX-design. Currently, 2 comprehensive BASIC linker sets are available ready to use on Opentrons in 96-well plates from Biolegio and more BASIC parts will be made available, enriching the design opportunities for new BASIC and DNA-BOT users^7,14^.

## Conclusion

Automated DNA assembly has largely been a reserve for well-funded institutions and Biofoundries. However, with the release of the Opentrons OT-2 pipetting robot, access to robotics is now within reach of most research groups. In this work we describe DNA-BOT, the implementation of BASIC DNA assembly on the Opentrons OT-2. Our dnabot package facilitates the generation of scripts to assemble and transform 96 BASIC constructs in a single run using up to 48 clip reactions. Importantly, all software and hardware required for this process are open-source and constructs can be generated for $1.50 worth of consumables and reagents. We utilised DNA-BOT to generate 88 constructs in parallel, each containing an operon expressing varying levels of GFP, RFP and BFP. The characterisation of these constructs highlighted the diversity of expression profiles DNA-BOT creates in this example application. Additionally, DNA-BOT offers high accuracy and efficiency, derived from the underlying BASIC DNA assembly method^5^. Convenient features at bench scale, they become a key advantage once automated DNA assembly is integrated into more automated Design-Built-Test-Learn workflows^19^.

## Supporting information

Storch_et_al_2019_SI

DNA_BOT_instructions_v1.0.0

## Author Contributions

M.S. and M. C. H. contributed equally to this work. M. S. and M. C. H. designed the methodology. M. C. H. performed the experiments and built the dnabot application. M.S and M.C.H. wrote the manuscript with contributions from G.S.B.

## Notes

The authors declare no competing financial interest.

## Notes

https://www.basic-assembly.org/

https://github.com/BASIC-DNA-ASSEMBLY/dnabot

