## Supplementary material for "DNA-BOT: A low-cost, automated DNA assembly platform for synthetic biology": Storch_et_al_2019_SI

### Materials and methods

#### BASIC DNA parts and availability

DNA parts from Table S1 with exception of BASIC_SEVA_37_CmR-p15A_v1.0 were cloned into a BsaI-site-free, Amp-pUC storage vector. The resulting plasmids and BASIC_SEVA_37_CmR-p15A_v1.0 were prepared at midiprep scale using GenScript®'s Plasmid DNA Prep Service and diluted to 200 ng/µL, ready to use in clip reactions.

A control plasmid containing only the backbone was derived from BASIC_SEVA_37_CmR-p15A_v1.0: BsaI digested plasmid was enzymatically blunted, self-ligated, transformed in DH5alpha and a correct clone identified via Sanger sequencing.

All DNA parts are available upon request.

#### DNA-BOT

A csv file describing each of the 88 constructs was generated along with csv files describing the Biolegio BASIC linker set (BBP-19100) standard linkers and the BASIC DNA parts required[1,2]. As described in DNA-BOT_instructions_v1.0.0 (Supporting Information), these csv files were used to generate four Opentrons OT-2 scripts. The 4$^{th}$ script was modified to transform and spot control plasmid and no plasmid controls in wells A12 – H12. Furthermore, a 5$^{th}$ script was generated separately to spot 10 µL of each transformation reaction. These scripts were implemented as described in DNA-BOT_instructions_v1.0.0 using LB-agar plates supplemented with 25 µg/mL chloramphenicol. All scripts are available at https://github.com/BASIC-DNA-ASSEMBLY/dnabot.

#### Flow cytometry

Individual colonies were picked from agar plates generated by DNA-BOT and 200 µL LB medium supplemented with 25 µg/mL chloramphenicol inoculated in 96-well plates. Cultures were incubated overnight, shaking 600 rpm at 30°C. Overnight cultures were diluted 200 times into 100 µL LB supplemented with 25 µg/mL chloramphenicol. Cultures were grown shaking at 30°C for 6 hours and 2 µL off-sampled into 200 µL Phosphate Buffer Saline supplemented with 2 mg/mL kanamycin. Samples were analysed for GFP, BFP and RFP fluorescence using a BD Fortessa flow cytometer with all samples gated using the same forward and side scatter settings. Data was analysed using FlowJo_V10 and subsequently processed as described in the main text.

### Supplementary figures

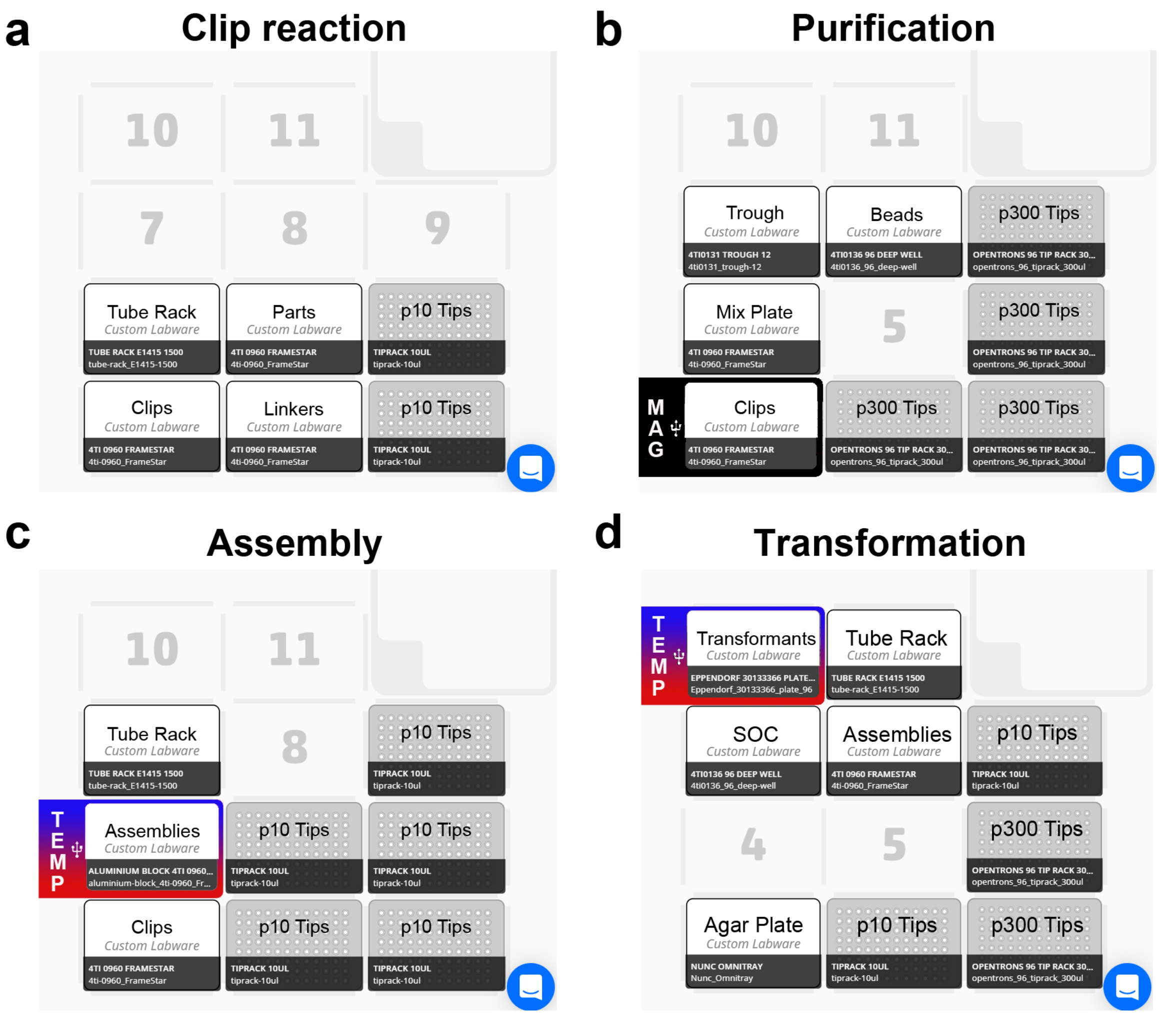

**Figure S1**. Deck layout used during generation of 88 constructs using DNA BOT (a) Script 1: Clip reaction, (b) Script 2: Purification, (c) Script 3: Assembly and (d) Script 4: Transformation.

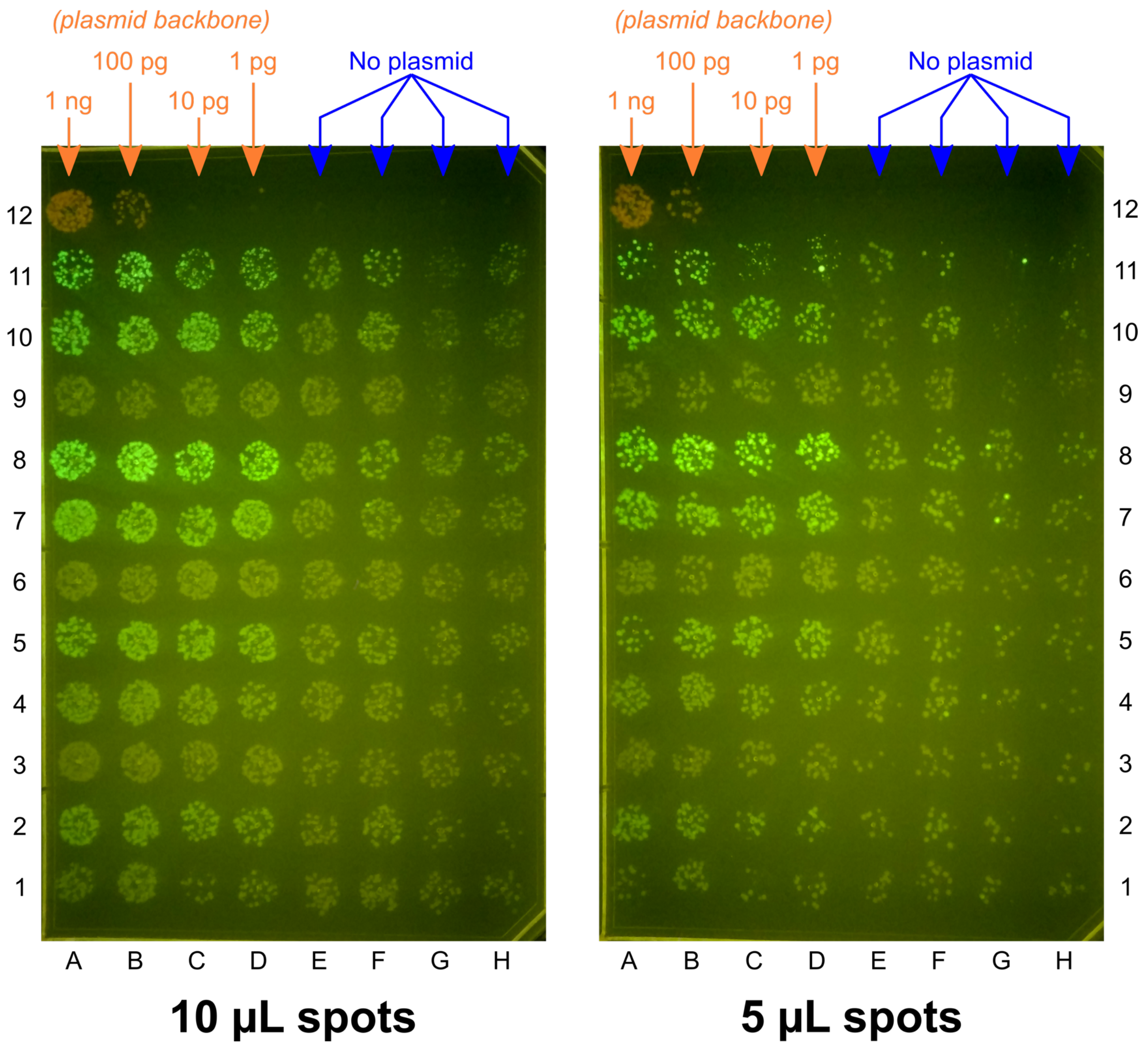

**Figure S2**. Agar plates imaged for GFP on a Safe Imager™ 2.0 Blue Light Transilluminator. Plates were spotted with 10 or 5 μL of each transformation reaction. Corresponding well identities can be inferred by the surrounding grid of letters and numbers. Cells spotted on positions A12 – H12, were transformed with 1 ng, 100 pg, 10 pg, 1 pg of BASIC_SEVA_37_CmR-p15A_v1.0 (plasmid backbone with mScarlet counter selection cassette) or with plasmid-free $H_2O$ (no plasmid), respectively. (Materials and methods).

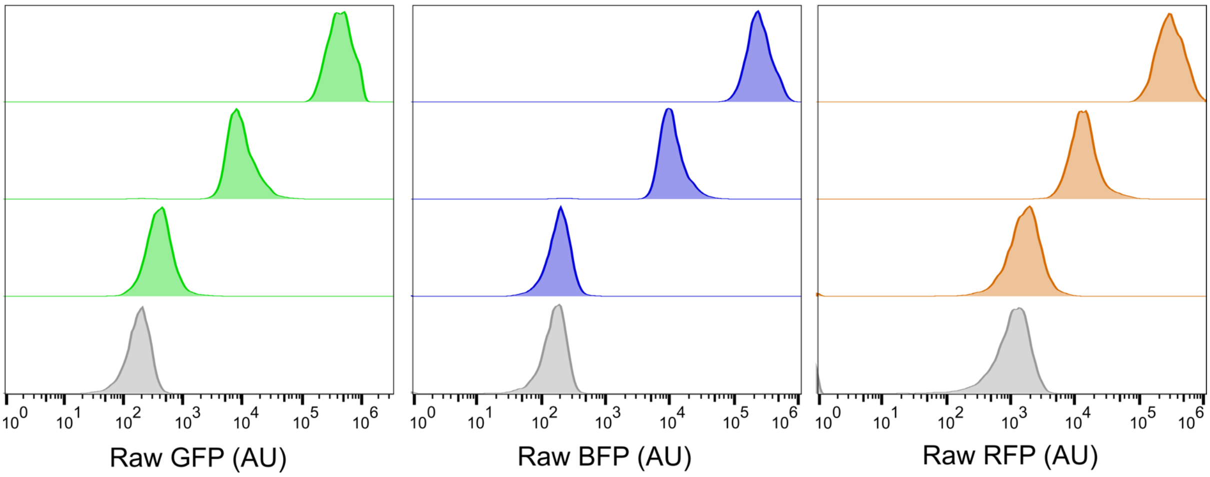

**Figure S3**. Flow cytometry histograms illustrating dynamic range in fluorescent reporter expression strength. Negative control cells with empty backbone are shown in grey and examples for low, medium and high expression phenotypes from the 88 construct library are shown in green (GFP), blue (BFP) and orange (RFP). Specifically, data for the following constructs are shown in order of low, medium and high expression, respectively: GFP: G3, H5 and C11. BFP expression: E2, H1 and H10. RFP: A1, F4 and H10.

### Supplementary tables

##### Table S1: DNA Sequences for BASIC parts used in this work

BASIC prefix (TCTGGTGGGTCTCTGTCC) and suffix (GGCTCGGGAGACCTATCG) sequences, along with the mScarlet coding sequence (magenta) are highlighted.

| ID | SynBioHub URI |
|---|---|
| BASIC_L3S2P21_J23105_RiboJ | TCTGGTGGGTCTCTGTCCCTCGGTACCAAATTCCAGAAAAGAGGCCTCCCGAAAGGGGGGCCTTTTTTCGTTTTGGTCCGTGCCTACTCTGGAAAATCTTTTACGGCTAGCTCAGTCCTAGGTACTATGCTAGCAGCTGTCACCGGATGTGCTTTCCGGTCTGATGAGTCCGTGAGGACGAAACAGCCTCTACAAATAATTTTGTTTAAGGCTCGGGAGACCTATCG |
| BASIC_L3S2P21_J23106_RiboJ | TCTGGTGGGTCTCTGTCCCTCGGTACCAAATTCCAGAAAAGAGGCCTCCCGAAAGGGGGGCCTTTTTTCGTTTTGGTCCGTGCCTACTCTGGAAAATCTTTTACGGCTAGCTCAGTCCTAGGTATAGTGCTAGCAGCTGTCACCGGATGTGCTTTCCGGTCTGATGAGTCCGTGAGGACGAAACAGCCTCTACAAATAATTTTGTTTAAGGCTCGGGAGACCTATCG |
| BASIC_L3S2P21_J23101_RiboJ | TCTGGTGGGTCTCTGTCCCTCGGTACCAAATTCCAGAAAAGAGGCCTCCCGAAAGGGGGGCCTTTTTTCGTTTTGGTCCGTGCCTACTCTGGAAAATCTTTTACAGCTAGCTCAGTCCTAGGTATTATGCTAGCAGCTGTCACCGGATGTGCTTTCCGGTCTGATGAGTCCGTGAGGACGAAACAGCCTCTACAAATAATTTTGTTTAAGGCTCGGGAGACCTATCG |
| BASIC_L3S2P21_J23104_RiboJ | TCTGGTGGGTCTCTGTCCCTCGGTACCAAATTCCAGAAAAGAGGCCTCCCGAAAGGGGGGCCTTTTTTCGTTTTGGTCCGTGCCTACTCTGGAAAATCTTTGACAGCTAGCTCAGTCCTAGGTATTGTGCTAGCAGCTGTCACCGGATGTGCTTTCCGGTCTGATGAGTCCGTGAGGACGAAACAGCCTCTACAAATAATTTTGTTTAAGGCTCGGGAGACCTATCG |
| BASIC_sfGFP_ORF_v1.0 | TCTGGTGGGTCTCTGTCC**ATG**CGTAAAGGCGAAGAACTGTTCACGGGCGTAGTTCCGATTCTGGTCGAGCTGGACGGCGATGTGAACGGTCATAAGTTTAGCGTTCGCGGTGAAGGTGAGGGCGACGCGACCAACGGCAAACTGACCCTGAAGTTCATCTGCACCACCGGTAAACTGCCGGTGCCTTGGCCGACCTTGGTGACGACGTTGACGTATGGCGTGCAGTGTTTTGCGCGTTATCCGGACCACATGAAACAACACGATTTCTTCAAATCTGCGATGCCGGAGGGTTACGTCCAGGAGCGTACCATTTCCTTCAAGGATGATGGCACTTACAAAACTCGCGCAGAGGTTAAGTTTGAAGGTGACACGCTGGTCAATCGTATCGAATTGAAGGGTATCGACTTTAAAGAGGATGGTAACATTCTGGGCCATAAACTGGAGTATAACTTCAACAGCCATAATGTTTACATTACGGCAGACAAGCAAAAGAACGGCATCAAGGCCAATTTCAAGATTCGCCACAATGTTGAGGACGGTAGCGTCCAACTGGCCGACCATTACCAGCAGAACACCCCAATTGGTGACGGTCCGGTTTTGCTGCCGGATAATCACTATCTGAGCACCCAAAGCGTGCTGAGCAAAGATCCGAACGAAAAACGTGATCACATGGTCCTGCTGGAATTTGTGACCGCTGCGGGCATCACCCACGGTATGGACGAGCTGTATAAGCGTCCGTAAGGCTCGGGAGACCTATCG |
| BASIC_RFP_ORF_v1.0 *(mCherry)* | TCTGGTGGGTCTCTGTCC**ATG**GTGAGCAAGGGCGAGGAGGATAACATGGCCATCATCAAGGAGTTCATGCGCTTCAAGGTGCACATGGAGGGCTCCGTGAACGGCCACGAGTTCGAGATCGAGGGCGAGGGCGAGGGCCGCCCCTACGAGGGCACCCAGACCGCCAAGCTGAAGGTGACCAAGGGTGGCCCCCTGCCCTTCGCCTGGGACATCCTGTCCCCTCAGTTCATGTACGGCTCCAAGGCCTACGTGAAGCACCCCGCCGACATCCCCGACTACTTGAAGCTGTCCTTCCCCGAGGGCTTCAAGTGGGAGCGCGTGATGAACTTCGAGGACGGCGGCGTGGTGACCGTGACCCAGGACTCCTCCTTGCAGGACGGCGAGTTCATCTACAAGGTGAAGCTGCGCGGCACCAACTTCCCCTCCGACGGCCCCGTAATGCAGAAGAAGACCATGGGCTGGGAGGCCTCCTCCGAGCGGATGTACCCCGAGGACGGCGCCCTGAAGGGCGAGATCAAGCAGAGGCTGAAGCTGAAGGACGGCGGCCACTACGACGCTGAGGTCAAGACCACCTACAAGGCCAAGAAGCCCGTGCAGCTGCCCGGCGCCTACAACGTCAACATCAAGTTGGACATCACCTCCCACAACGAGGACTACACCATCGTGGAACAGTACGAACGCGCCGAGGGCCGCCACTCCACCGGCGGCATGGACGAGCTGTACAAGTAAGGCTCGGGAGACCTATCG |
| BASIC_BFP_ORF_v1.0 | TCTGGTGGGTCTCTGTCC**ATG**TCCGAGTTGATCAAAGAGAACATGCATATGAAATTATATATGGAAGGCACTGTAGATAATCATCATTTTAAATGTACGTCGGAAGGCGAAGGTAAACCATATGAAGGTACGCAGACGATGCGCATCAAGGTGGTGGAGGGCGGTCCGCTGCCATTCGCTTTCGATATTTTAGCCACGAGCTTCCTCTACGGTTCTAAAACTTTCATCAATCACACGCAGGGTATTCCGGACTTCTTTAAACAGTCGTTCCCGGAGGGTTTCAC |

| | |
|---|---|
| | CTGGGAACGCGTTACCACGTATGAAGATGGTGGTGTGCTTACGGCAACGCAGGACACGAGCCTTCAGGATG<br>GGTGTTTGATTTACAACGTGAAAATTCGTGGTGTGAACTTCACGTCTAACGGCCCGGTGATGCAGAAAAAA<br>ACACTGGGTTGGGAAGCCTTTACCGAAACCCTGTATCCGGCGGACGGTGGCCTGGAAGGCCGTAATGATAT<br>GGCCTTGAAATTAGTCGGCGGTTCACACCTGATCGCGAACGCGAAAACAACCTATCGTAGTAAAAAACCAG<br>CCAAAAACCTGAAAATGCCGGGCGTCTACTACGTAGACTACCGTCTGGAGCGCATTAAAGAGGCGAATAAT<br>GAAACCTATGTCGAGCAGCACGAAGTTGCGGTTGCACGCTATTGCGATCTGCCCAGCAAACTGGGCCACAA<br>GCTTAATGGTAGCTAAGGCTCGGGAGACCTATCG |
| BASIC_SEVA_37_CmR-p15A_v1.0 | TCTGGTGGGTCTCTGTCCACTAGTCTTGGACTCCTGTTGATAGATCCAGTAATGACCTCAGAACTCCATCTGGATTTGTTCAGAACGCTCGGTTGCCGCCGGGCGTTTTTTAT<br>TGGTGAGAATCCAGGGGTCCCCAATAATTACGATTTAAATTGGCGAAAATGAGACGTTGATCGGCACGTAAGAGGTTCCAACTTTCACCATAATGAAATAAGATCACTACCGG<br>GCGCTGAGAGATCCCCTCATAATTTCCCCAAAGCGTAACCATGTGTGAATAAATTTTGAGCTAGTAGGGTTGCAGCCACGAGTAAGTCTTCCCTTGTTATTGTGTAGCCAGAA<br>TGCCGCAAAACTTCCATGCCTAAGCGAACTGTTGAGAGTACGTTTCGATTTCTGACTGTGTTAGCCTGGAAGTGCTTGTCCCAACCTTGTTTCTGAGCATGAACGCCCGCAAG<br>CCAACATGTTAGTTGAAGCATCAGGGCGATTAGCAGCATGATATCAAAACGCTCTGAGCTGCTCGTTCGGCTATGGCGTAGGCCTAGTCCGTAGGCAGGACTTTTCAAGTCTC<br>GGAAGGTTTCTTCAATCTGCATTCGCTTCGAATAGATATTAACAAGTTGTTTGGGTGTTCGAATTTCAACAGGTAAGTTAGTTGCTAGAACCCATGGCTCCTTTGCCGACGCT<br>GAGTAGATTTTAGGTGACGGGTGGTGACAATGAGTCCGTGTCGAGCGCTGATTTTTTCGGCCTTTAGAGCGAGATTTATACAATAGAATTTGGCATGAGATTGGATTGCTTTT<br>AGTCAGCCTCTTATAGCCTAAAGTCTTTGAGTGACTAGATGACATATCATGTAAGTTGCTGATAGGTTTCCAGTTTTCCGCTCCTAGGTCTGCATATTGTACTTTTCCTCTTA<br>CTCGACTTAACCAGTACCAACCCAGCTTCTCAACGGATTTATACCATGGCACTTTAAAGCCAGCATCACTGACAATGAGCGGTGTGGTGTTACTCGGTAGAATGCTCGCAAGG<br>TCGGCTAGAAATTGGTCATGAGCTTTCTTTGAACATTGCTCTGAAAGCGGGAACGCTTTCTCATAAAGAGTAACAGAACGACCGTGTAGTGCGACTGAAGCTCGCAATACCAT<br>AAGTCGTTTTTGCTCACGAATATCAGACCAGTCAACAAGTACAATGGGCATCGTATTGCCCGAACAGATAAAGCTAGCATGCCAACGGTATACAGCGAGTCGCTCTTTGTGGA<br>GGTGACGATTACCTAACAATCGGTCGATTCGTTTGATGTTATGTTTTGTTCTCGCTTTGGTTGGCAGGTTACGGCCAAGTTCGGTAAGAGTGAGAGTTTTACAGTCAAGTAAT<br>GCGTGGCAAGCCAACGTTAAGCTGTTGAGTCGTTTTAAGTGTAATTCGGGGCAGAATTGGTAAAGAGAGTCGTGTAAAATATCGAGTTCGCACATCTTGTTGTCTGATTATTG<br>ATTTTTCGCGAAACCATTTGATCATATGACAAGATGTGTATCCACCTTAACTTAATGATTTTTACCAAAATCATTAGGGGATTCATCAGTACCGGGCGTATTTTTTGAGTTAT<br>CGAGATTTTCAGGAGCTAAGGAAGCTAAAATGGAGAAAAAAATCACTGGATATACCACCGTTGATATATCCCAATGGCATCGTAAAGAACATTTTGAGGCATTTCAGTCAGTT<br>GCTCAATGTACCTATAACCAGACCGTTCAGCTGGATATTACGGCCTTTTTAAAGACCGTAAAGAAAAATAAGCACAAGTTTTATCCGGCCTTTATTCACATTCTTGCCCGCCT<br>GATGAATGCTCATCCGGAATTTCGTATGGCAATGAAAGACGGTGAGCTGGTGATATGGGATAGTGTTCACCCTTGTTACACCGTTTTCCATGAGCAAACTGAAACGTTTTCAT<br>CGCTCTGGAGTGAATACCACGACGATTTCCGGCAGTTTCTACACATATATTCGCAAGATGTGGCGTGTTACGGTGAAAACCTGGCCTATTTCCCTAAAGGGTTTATTGAGAAT<br>ATGTTTTTCGTCTCAGCCAATCCCTGGGTGAGTTTCACCAGTTTTGATTTAAACGTGGCCAATATGGACAACTTCTTCGCCCCCGTTTTCACCATGGGCAAATATTATACGCA<br>AGGCGACAAGGTGCTGATGCCGCTGGCGATTCAGGTTCATCATGCCGTTTGTGATGGCTTCCATGTCGGCAGAATGCTTAATGAATTACAACAGTACTGCGATGAGTGGCAGG<br>GCGGGGCGTAATTTGACTTTTGTCGGCTCGACCCACGACTATTGACTGCTCTGAGAAAGTTGATTGTTACGATTAGTCCGGCCGGCCCTAGAAATATTTTATCTGATTAATAA<br>GATGATCTTCTTGAGATCGTTTTGGTCTGCGCGTAATCTCTTGCTCTGAAAACGAAAAAACCGCCTTGCAGGGCGGTTTTTCGAAGGTTCTCTGAGCTACCAACTCTTTGAAC<br>CGAGGTAACTGGCTTGGAGGAGCGCAGTCACCAAAACTTGTCCTTTCAGTTTAGCCTTAACCGGCGCATGACTTCAAGACTAACTCCTCTAAATCAATTACCAGTGGCTGCTG<br>CCAGTGGTGCTTTTGCATGTCTTTCCGGGTTGGACTCAAGACGATAGTTACCGGATAAGGCGCAGCGGTCGGACTGAACGGGGGGTTCGTGCATACAGTCCAGCTTGGAGCGA<br>ACTGCCTACCCGGAACTGAGTGTCAGGCGTGGAATGAGACAAACGCGGCCATAACAGCGGAATGACACCGGTAAACCGAAAGGCAGGAACAGGAGAGCGCACGAGGGAGCCGC<br>CAGGGGGAAACGCCTGGTATCTTTATAGTCCTGTCGGGTTTCGCCACCACTGATTTGAGCGTCAGATTTCGTGATGCTTGTCAGGGGGGCGGAGCCTATGGAAAAACGGCTTT<br>TGCCGCGGCCCTCTCACTTCCCTGTTAAGTATCTTCCTGGCATCTTCCAGGAAATCTCCGCCCCGTTCGTAAGCCATTTCCGCTCGCCGCAGTCGAACGACCGAGCGTAGCGA<br>GTCAGTGAGCGAGGAAGCGGAATATATCCGGCGCGCCCAGCTGTCTAGGGCGGCGGATTTGTCCTACTCAGGAGAGCGTTCACCGACAAACAACAGATAAAACGAAAGGCCCA<br>GTCTTTCGACTGAGCCTTTCGTTTTATTTGATGCCTTTAATTAAGGCTCGGGAGACCTATCGGTAATAACAGTCCAATCTGGTGTAACTTCGGAATCGTCCTTGACAGCTAGC<br>TCAGTCCTAGGTATAATGCTAGCTACTAGTGAAAGAGGAGAAATACTAGTATGGTTAGCAAAGGCGAGGCGGTTATCAAGGAGTTTATGCGTTTTAAGGTTCACATGGAGGGT<br>AGCATGAATGGTCACGAGTTCGAGATCGAGGGTGAAGGCGAGGGTCGTCCGTACGAAGGCACCCAGACCGCGAAGCTGAAAGTGACCAAGGGTGGCCCGCTGCCGTTCAGCTG<br>GGACATCCTGAGCCCGCAGTTCATGTATGGCAGCCGTGCGTTTACCAAACACCCGGCGGACATTCCGGATTACTATAAGCAAAGCTTCCCGGAAGGTTTTAAATGGGAGCGTG<br>TTATGAACTTCGAAGATGGTGGCGCGGTGACCGTTACCCAGGACACCAGCCTGGAGGATGGCACCCTGATTTACAAGGTGAAACTGCGTGGCACCAACTTTCCGCCGGATGGT<br>CCGGTTATGCAGAAGAAAACGATGGGTTGGGAAGCGAGCACCGAGCGTCTGTATCCGGAAGATGGCGTGCTGAAGGGTGATATCAAAATGGCGCTGCGTCTGAAGGACGGTGG<br>CCGTTACCTGGCGGATTTTAAGACCACCTATAAAGCGAAGAAACCGGTGCAAATGCCGGGTGCGTACAACGTTGACCGTAAACTGGATATTACCAGCCACAACGAGGATTATA<br>CCGTGGTTGAGCAATATGAGCGTAGCGAGGGTCGCCACAGCACCGGCGGCATGGACGAACTGTATAAGGGATCCTAATAACGCTGATAGTGCTAGTGTAGATCGCTACTAGAG<br>CCAGGCATCAAATAAAACGAAAGGCTCAGTCGAAAGACTGGGCCTTTCGTTTTATCTGTTGTTTGTCGGTGAACGCTCTCTACTAGAGTCACACTGGCTCACCTTCGGGTGGG<br>CCTTTCTGCGTTTATATACTGGCTCGGGTAAGAACTCGCACTTCGTGGAAACACTATTA |

**Table S2: DNA Sequences for BASIC linkers used in this work**

UTR-RBS linkers were used to build the operons and methylated linkers LMP & LMS were used to connect the expression cassette with the backbone. The 3 Biobrick RBS sequences (underlined) encoded on the UTR-RBS linkers were in order from low to high translation strength: BBa_B0033 (RBS1), BBa_B0064 (RBS2), BBa_B0034 (RBS3). Central uppercase sequence sections indicate the homology regions of the single stranded 21 base overhangs, which bring Clips together during final assembly.

| | |
|---|---|
| UTR1-RBS1 | ctcgttgaacaccgtcTCAGGTAAGTATCAGTTGTAAatcacacaggactagtcc |
| UTR1-RBS2 | ctcgttgaacaccgtcTCAGGTAAGTATCAGTTGTAAaaagaggggaaatagtcc |
| UTR1-RBS3 | ctcgttgaacaccgtcTCAGGTAAGTATCAGTTGTAAaaagaggagaaatagtcc |
| UTR2-RBS1 | ctcgtgttactattggCTGAGATAAGGGTAGCAGAAAatcacacaggactagtcc |
| UTR2-RBS3 | ctcgtgttactattggCTGAGATAAGGGTAGCAGAAAaaagaggagaaatagtcc |
| UTR3-RBS1 | ctcggtatctcgtggtCTGACGGTAAAATCTATTGTAatcacacaggactagtcc |
| UTR3-RBS3 | ctcggtatctcgtggtCTGACGGTAAAATCTATTGTAaaagaggagaaatagtcc |
| LMP | ctcgggtaagaactcgCACTTCGTGGAAACACTATTAtctggtgggtctctgtcc |
| LMS | ctcgggagacctatcgGTAATAACAGTCCAATCTGGTGTaacttcggaatcgtcc |

**Table S3: DNA-BOT running costs per construct**

Costing assumes constructs are composed of 5 parts and each purified clip is used in 15 assemblies, representative for the 88 constructs. Costs for foil seals, linkers, trough, microcentrifuge tubes, assembly buffer, ethanol and water were considered neglectable. Unit prices correct as of October 2019 and values give to 2 DP.

| **Item** | **Units** | **Units per construct** | **Unit price (USD)** | **Cost (USD)** |
|---|---|---|---|---|
| Opentrons p10 tips | tips | 9.33 | 0.03 | 0.27 |
| Framestar 96-well Rigid and Skirted PCR Plates | plates | 0.02 | 3.69 | 0.06 |
| Promega T4 DNA Ligase | µL | 0.17 | 1.18 | 0.20 |
| NEB BsaI-HF®v2 | µL | 0.33 | 1.11 | 0.37 |
| Opentrons p300 tips | tips | 5.00 | 0.03 | 0.14 |
| Brooks Life Sciences 96 Square Deep Well Storage Microplate | plates | 0.01 | 3.69 | 0.04 |
| Beckman Coulter™ Agencourt AMPure XP SPRI paramagnetic beads | mL | 0.02 | 16.62 | 0.30 |
| *NEB® 5-alpha Competent E. coli, 96 well plate* | *plate* | *0.01* | *380.12* | *3.96* |
| Thermo ScientificTM Nunc™ OmniTray™ Single-Well Plate | plate | 0.01 | 3.58 | 0.04 |
| **Total** | | | | **5.37** |
| **Total – w/o cells** | | | | **1.41** |

**Table S4: Hands-on time required for manual BASIC assembly and DNA-BOT.**

Calculated for the implementation of 48 clip reactions, plus the assembly and transformation of 88 constructs. The $Q_{time}$ metric[3] has been calculated from these two values.

| **Step** | **Manual protocol (mins)** | **Automated protocol (mins)** |
|---|---|---|
| Clip reactions (48) | 60 | 15 |
| Purification | 30 | 25 |
| Assembly | 90 | 15 |
| Transformation | 100 | 35 |
| **Total** | **280** | **90** |
| **$Q_{time}$ = 3.11** | | |

### Bibliography

(1) Biolegio BASIC linkers https://www.biolegio.com/products-services/basic/.

(2) BASIC Assembly website www.basic-assembly.org.

(3) Walsh, D. I.; Pavan, M.; Ortiz, L.; Wick, S.; Bobrow, J.; Guido, N. J.; Leinicke, S.; Fu, D.; Pandit, S.; Qin, L.; et al. Standardizing Automated DNA Assembly: Best Practices, Metrics, and Protocols Using Robots. *SLAS Technol. Transl. Life Sci. Innov.* **2019**, 247263031882533. https://doi.org/10.1177/2472630318825335.
