## Supplementary material for "DNA-BOT: A low-cost, automated DNA assembly platform for synthetic biology": DNA_BOT_instructions_v1.0.0

### Instructions for DNA-BOT (v1.0.0)

#### About

This document is intended to guide users through DNA assembly using BASIC on OpenTrons (DNA-BOT).

#### Materials

Below we list the materials previously used to implement DNA-BOT. We recommend starting with these consumables, however certain materials like 96-well plates, microcentrifuge tubes etc. may be altered. Labware needs to be defined carefully when working on the Opentrons OT-2 and changes to labware require the user to update template scripts[1] or those executed during DNA-BOT runs. We further advise users to follow general opentrons guidelines when utilising labware.

Software:

- Opentrons OT-2 Run App (Version 3.10.3 or later).
- dnabot GitHub repository.
- Python 3 and packages specified in requirements.txt.

Hardware:

- Opentrons OT-2.
- Opentrons P10 Single-Channel Electronic Pipette.
- Opentrons P300 8-Channel Electronic Pipette.
- Opentrons Magnetic Module.
- Opentrons Temperature Module.
- Opentrons 4-in-1 Tube Rack Set.
- Opentrons Aluminum Block Set.
- Thermocycler compatible with 96-well plates.
- Microplate centrifuge.
- General laboratory equipment e.g. pipettes.

Consumables & reagents:

| **Item** | **Cat no./URL** |
| --- | --- |
| Opentrons 10µL tips | https://shop.opentrons.com/products/opentrons-10ul-tips |
| Opentrons 300µL tips | https://shop.opentrons.com/collections/opentrons-tips/products/opentrons-300ul-tips |
| Framestar 96-well Rigid and Skirted PCR Plates | 4ti-0960/RIG |
| Brooks Life Sciences Reservoir Plate, 21 mL channels | 4ti-0131 |
| Brooks Life Sciences 96 Square Deep Well Storage Microplate, 2.2 mL wells, U shaped bases | 4ti-0136 |
| Brooks Life Sciences PCR Foil Seals | 4ti-0550 |

[1]Note changes to template scripts will apply to all implementations of the dnabot app.

| | |
|---|---|
| Thermo Scientific™ Nunc™ OmniTray™ Single-Well Plate, Non-treated | 242811 |
| 1.5 mL STARLAB Crystal Clear Microcentrifuge tubes | E1415-1500 |
| [optional] Brooks Life Sciences 96 Square Well Sealing Cap Mat | 4ti-0137 |
| Biolegio BASIC linker set | BBP-19100 https://www.biolegio.com/products-services/basic/ |
| NEB BsaI-HF®v2 | R3733 |
| Promega T4 DNA Ligase | M1801 |
| 10x Assembly Buffer: 0.2 M Tris:HCl (pH 8.0), 0.1 M $MgCl_2$, 0.5 M KCl | N/A – generate from component materials (Sambrook and Russell 2001). |
| Beckman Coulter™ Agencourt AMPure XP SPRI paramagnetic beads | A6388 |
| NEB® 5-alpha Competent E. coli, 96 well plate | C2987P |

Additional reagents:

- ddH2O.
- Absolute ethanol.
- SOC media.
- LB-Agar + antibiotic e.g. chloramphenicol.

#### Methodology

##### OT-2 preparation

Follow the Opentrons guidelines for setting up the OT-2 before executing any protocols. 3rd-party labware not given in the default Opentrons OT-2 Labware Database but listed in the Materials section can be added to user OT-2 labware databases by running the add_labware.ot2.py script.

##### Construct and part/linker csv files

At least, two csv files are required to run the dnabot app, generating python scripts for the BASIC assembly on Opentrons OT-2. The first csv file contains the details of constructs to be assembled. Additional csv files are required for each 96-well plate containing BASIC parts and/or BASIC linkers that are used during clip reactions.

constructs_temp.csv provides a template to describe constructs for new DNA-BOT runs. As an example, storch_et_al_cons.csv describes constructs previously assembled. **CONSTRUCTS MUST BE LISTED CONSECUTIVELY, WITH THE FIRST CONSTRUCT BEING ASSEMBLED IN WELL A1**. Each construct encompasses a single row of the csv file and is composed of alternating BASIC linkers and BASIC DNA parts. Each linker sequence is the result of annealed prefix and suffix linker sections within the final construct. For instance, "LMS" refers to the sequence formed when "LMS-S" and "LMS-P" linkers anneal during the assembly step of BASIC[2]. Any linker supplied by Biolegio can be used to design constructs[3].

parts_linkers_temp.csv provides a template to describe plates containing parts or linkers, used during clip reactions. Additionally, BIOLEGIO_BASIC_STD_SET.csv provides an example of a linker plate previously used to assemble constructs. The first row of the csv file details the csv file header. The "Part/linker" column provides the ID of the part or linker. These exact ID values have to be used in the construct csv file, with prefix and suffix linker IDs extended by "-P" and "-S", respectively. Parts and linkers not used to assemble constructs listed in the construct csv file are ignored by the dnabot app during implementation. The "Well" column provides the corresponding well of the part or linker. No restrictions apply to well order and adding a part present in another well ensures only the latter is used. This enables new aliquots of parts to be added to the plate in separate wells without having to remove the previous instance from the csv file. The "Part concentration (ng/uL)" column provides the corresponding concentration of DNA parts. 2.5 nM final concentration of DNA part during clip reactions is optimal (Storch et al. 2015) and given clip reaction volume and typical DNA part sizes (0.1-2kb) in storage vectors, 200 ng DNA part is considered optimal by the dnabot app. BASIC DNA parts must be supplied at a concentration > 25 ng/μL. For concentrations > 200 ng/μL and if this value is left blank, 1 μL is pipetted.

---

[2] For additional understanding of the BASIC process, the original publication should be consulted (Storch et al. 2015).
[3] While additional, user-generated linkers are likely tolerated, they are currently discouraged.

#### Running the dnabot app

To acquire the dnabot app, clone the dnabot GitHub repository. Note, the dnabot package is not intended to be installed at the environment level and is instead a stand-alone application. The simplest way to start the dnabot app is to run the dnabot_app.py module from within the dnabot package using an appropriate Python environment. This will bring up a simple graphic user interface with instructions. Follow these instructions to generate the 4 scripts and metainformation required to execute the DNA-BOT workflow. Following app execution, scripts and metainformation are located in the directory containing the construct csv file.

#### Step 1: Clip reactions

##### *Preparation*

Prepare PCR plates containing DNA parts and linkers, ensuring no air gaps exist at the bottom of wells and that sufficient material is available. A dead volume of 10 – 15 µL for each part and linker is required to ensure complete liquid transfers. For consistent results, DNA parts should be well mixed prior to script execution.

Prepare the clip reaction MASTER_MIX in a microcentrifuge tube by combining the required materials specified in the "[construct csv file name]_clip_run_info.csv" file[4], located in the "metainformation" directory. Prepare an additional microcentrifuge tube containing dd$H_2O$.

##### *Script execution*

Execute the "1_clip.ot2.py" script using the Opentrons app and follow the instructions for calibration and running. An example deck layout is given below. Plates containing parts and linkers must be placed in deck positions as specified in the "[construct csv file name]_clip_run_info.csv" file. Additionally, place tubes containing master mix and water in positions A1 and A2 of the tube rack, respectively. A fresh PCR plate should be placed on deck position 1 for clip reactions.

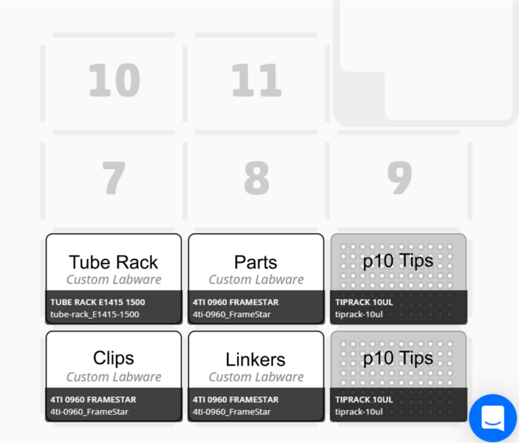

[4] Refer to storch_et_al_cons_clip_run_info.csv for a previously generated example.

***Thermocycler incubation***

Following completion of the “1_clip.ot2.py” script, remove the plate containing clip reactions from the OT-2. Seal the plate using a foil seal and transfer to a thermocycler. Conduct the following incubation:

| Temperature | Time | |
|---|---|---|
| 37°C | 2 min | **X 20 cycles** |
| 20°C | 1 min | |
| 37°C | 5 min | |
| 80°C | 20 min | |

#### Step 2: Purification

***Preparation***

Following clip reaction incubation, remove the foil seal from the plate containing raw clip reactions. Prepare a trough with $ddH_2O$ in well A1 and fresh 70 % EtOH[5] in the selected well[6]. Resuspend the SPRI beads and aliquot 0.75 mL into wells A1 – H1, inc. of a 96 deep-well plate[7].

***Script execution***

Execute the “2_purification.ot2.py” script using the app and follow the instructions for calibration and running. An example deck layout is given below. A 96 well plate, referred to as a “Mix Plate” is required for effective SPRI immobilisation.

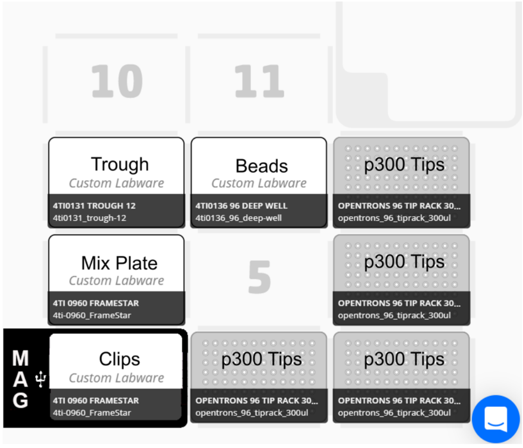

[5] Follow the SPRI bead manufacturers guidelines for the preparation of 70 % EtOH.
[6] If required, refer to the metainformation directory or “2_purification.ot2.py” script for the previously selected well.
[7] Beads can be stored within the 96 deep well plate using the corresponding seal mats. To resuspend within the plate simply invert several times before removing the seal. Ensure sufficient bead volume is available in each well before executing scripts.

#### Step 3: Assembly

##### *Preparation*

Optionally seal the plate containing purified clips with a foil seal and centrifuge to guarantee the absence of air gaps in wells. Air gaps in wells can cause liquid transfers during assemblies to fail.

Assemblies consisting of a different number of parts, require different dilutions of 10x Assembly Buffer (see table below). This functions to reduce the number of pipetting steps, saving time and resources. The trade-off is that multiple solutions must be generated when assembling constructs containing different part numbers. However, Assembly Buffer stores well at 4 °C meaning dilutions can be prepared in bulk (≥ 10 mL), aliquoted and stored until required (see table below).

Prepare the required dilutions of Assembly Buffer for the constructs being assembled in microcentrifuge tubes. Place the tube in the specified tube rack well (see table below).

| Number of parts to assemble | [Assembly Buffer] (fold) | 10x Assembly Buffer for 10 mL (mL) | ddH$_2$O for 10 mL (mL) | Suggested aliquot (mL) | Well in tube rack |
|---|---|---|---|---|---|
| 2 | 1.3 | 1.3 | 8.7 | 1.25 | A1 |
| 3 | 1.4 | 1.4 | 8.6 | 1.1 | A2 |
| 4 | 1.7 | 1.7 | 8.3 | 0.95 | A3 |
| 5 | 2 | 2 | 8 | 0.8 | A4 |
| 6 | 2.5 | 2.5 | 7.5 | 0.65 | A5 |

##### *Script execution*

Execute the “3_assembly.ot2.py” script using the app and follow the instructions for calibration and running. An example deck layout is given below. A fresh, 96-well PCR plate should be used for assemblies. This plate is placed on the aluminium 96-well block which in turn is placed on the Temperature Module.

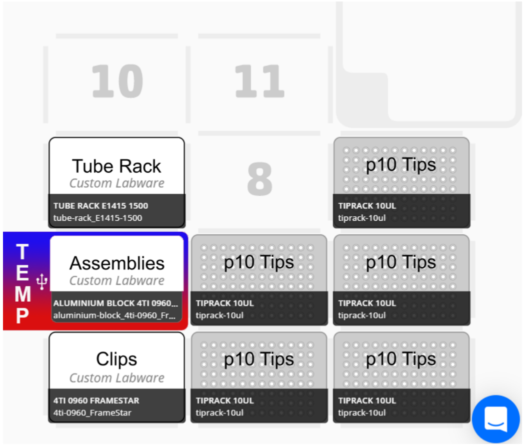

##### *Thermocycler incubation*

Following completion of the "3_assembly.ot2.py" script, remove the plate containing assemblies from the OT-2. Seal the plate using a foil seal and transfer to a thermocycler. Conduct the following incubation:

| Temperature | Time |
|---|---|
| 50°C | 45 min |
| 4°C | hold |

#### Step 4: Transformation

##### *Preparation*

Remove the plate containing the assemblies from the thermocycler and remove the foil seal.

Prepare a plate for SOC media by first sterilising a 96 deep-well plate and then filling each well of the selected column[8] with 2 mL sterile SOC media. Avoid touching neighbouring wells, enabling remaining columns to be used during subsequent runs.

Prepare the agar plate by pouring 40 mL LB-agar supplemented with the required concentration of antibiotic/s into a Nunc™ Omnitray™. Leave to set, and dry prior to use[9].

Place the aluminium 96-well block on ice or at 4 °C to chill. Place the plate containing competent cells on the chilled block, remove the foil and cover with the provided plate cover. Place the block with cells on ice until required.

##### *Script execution*

Execute the "4_transformation.ot2.py" script using the app and follow the initial instructions for pipette calibration. An example deck layout is given below. Continue the calibration protocol, calibrating pipette tips. To calibrate well A1 of the agar plate, first calibrate the x and y-axes using a 96-well PCR plate. Remove the PCR plate and reintroduce the agar plate to calibrate the z-axis, move the pipette tip until it just touches the agar. Continue with the instructions to calibrate the remaining labware. The tube rack requires a single empty microcentrifuge tube in well A1 for spotting waste. Once the plate containing cells has been calibrated, cover and store on ice until prompted to return[10].

Proceed by running the protocol. The script will first cool the temperature deck to 4 °C, before pausing, allowing users to place competent cells on the deck. After resuming, the OT-2 will mix assemblies with competent cells and incubate for 20 minutes. The user is then prompted to conduct a

---

[8] If required, refer to the metainformation directory or "4_transformation.ot2.py" script for the previously selected column of wells.

[9] To avoid spots fusing during spotting, agar plates must be well dried. Previously, plates have successfully been dried at 37 °C for 3 – 4 hours.

[10] The agar plate and SOC media can also be removed from the deck and covered until after heat shock, to reduce the risk of contamination.

heat shock. To achieve this, preheat a thermocycler to 42 °C and remove cells to ice. Heat shock at 42 °C for 15 seconds in the thermocycler and return to ice for 2 minutes. Transfer the cells back to the temperature deck and resume the run. No further pauses are implemented, allowing users to return after script completion.

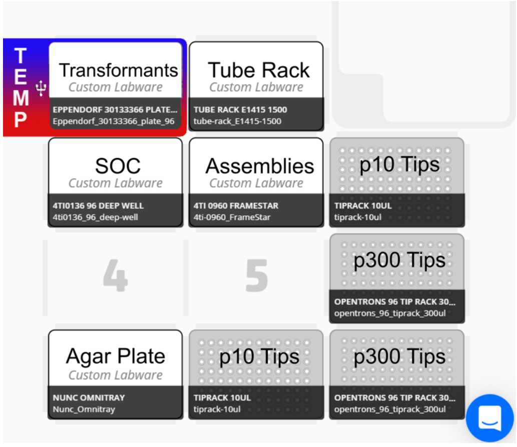

#### Colony generation and picking

Following spotting, incubate agar plates at 37 °C until colony formation and apply a foil seal to the plate containing transformation reactions. Transformation reactions can be stored as backup at 4 °C for several days.

To isolate single colonies for downstream applications users may be required to conduct further manual steps. When no colonies form following spotting, the remaining transformation reaction should be plated on an agar plate (Sambrook and Russell 2001). This facilitates the screening of larger volumes of transformation reaction for the presence of successful transformants. In addition to observing no colonies, it is possible that colony density is too high for single-colony isolation. This is likely to be the case for assemblies composed of only 2 or 3 parts, where the transformation efficiency is higher. To overcome this, spots with high colony densities should be streaked-out (Sambrook and Russell 2001).

#### Bibliography

Sambrook, Joseph, and David Russell. 2001. *Molecular Cloning A Laboratory Manual*. 3rd ed. New York: Cold Spring Harbor Laboratory Press.

Storch, Marko, Arturo Casini, Ben Mackrow, Toni Fleming, Harry Trewhitt, Tom Ellis, and Geoff S. Baldwin. 2015. “BASIC: A New Biopart Assembly Standard for Idempotent Cloning Provides Accurate, Single-Tier DNA Assembly for Synthetic Biology.” *ACS Synthetic Biology* 4 (7): 781–87. https://doi.org/10.1021/sb500356d.
